# Increased cortical inhibition immediately following brief motor memory reactivation supports reconsolidation and overnight offline learning gains

**DOI:** 10.1101/2023.02.21.529358

**Authors:** Tamir Eisenstein, Edna Furman-Haran, Assaf Tal

## Abstract

Practicing motor skills stabilizes and strengthens motor memories by repeatedly reactivating and reconsolidating them. The conventional view, by which a repetitive practice is required for substantially improving skill performance, has been recently challenged by behavioral experiments, in which even brief reactivations of the motor memory have led to significant improvements in skill performance. However, the mechanisms which facilitate brief reactivation-induce skill improvements remain elusive. While initial memory consolidation has been repeatedly associated with increased neural excitation and dis-inhibition, reconsolidation has been shown to involve a poorly-understood mixture of both excitatory and inhibitory alterations.

Here, we followed a three-day reactivation-reconsolidation framework, to examine whether the excitatory/inhibitory mechanisms which underlie brief reactivation and repetitive practice differ. Healthy volunteers practiced a motor sequence learning task using either brief reactivation or repetitive practice and were assessed using ultra-high field (7T) magnetic resonance spectroscopy at the primary motor cortex (M1). We found that increased inhibition (GABA concentrations) and decreased excitation/inhibition (glutamate/GABA ratios) immediately following the brief reactivation were associated with overnight offline performance gains. These gains were on-par with those exhibited following repetitive practice, where no correlations with inhibitory or excitatory changes were observed. Our findings suggest that brief reactivation and repetitive practice depend on fundamentally different neural mechanisms, and that early inhibition – and not excitation – is particularly important in supporting the learning gains exhibited by brief reactivation.

## Introduction

The acquisition of new motor skills is ubiquitous in our daily lives, playing a crucial role in learning how to grasp a spoon as babies, playing a new musical instrument as adults, and recovering function following acute brain injury. While traditionally it has been accepted that ‘practice makes perfect’ when it comes to learning and further refining and improving the performance of a motor skill over time^1^, recent studies across different modalities of procedural learning, including the motor domain, have started to question this dogma^2–4^. According to those findings, briefly reactivating a memory of a previously learned skill promotes improved skill performance and may even be as beneficial as practicing it extensively/repetitively^2^. While the learning-induced potential of brief reactivation may hold far reaching implications in clinical and rehabilitative settings such as following stroke and brain injury, this phenomenon has been mainly demonstrated behaviourally, and the neural processes which may support it are not well understood. In particular, it is unclear whether the mechanisms underlying brief reactivation-induced learning somehow differ from those of more conventional motor learning approaches.

The effectiveness of a brief reactivation in promoting further skill learning is potentially linked to its ability to modify the existing motor memory trace by initiating *reconsolidation* processes^5–7^. Following its reactivation, a memory trace has been shown to become labile again and prone to modifications^6,8–11^, and reconsolidation is needed in order to restabilize it (i.e., making the memory once again resistant to interference) and promote skill improvements that develop after the practice session (i.e., offline learning gains). Neural inhibition and excitation (E-I) have been suggested to play a key role in the acquisition and consolidation of new motor skills in both animals and humans^12–15^; we have therefore hypothesized they also play a fundamental role during the reconsolidation initiated by a brief reactivation. Changes in glutamate (Glu) and γ-aminobutyric acid (GABA), the main excitatory and inhibitory neurotransmitters in the brain, have been documented in the primary motor cortex (M1) both during^12,13^ and following^14,15^ the initial acquisition of motor skills. Motor learning in humans has been previously characterized by GABAergic-mediated cortical disinhibition during learning^12,13^. Furthermore, recent findings in humans have demonstrated that the initial consolidation of motor memories may also depend on increased cortical excitation and GABAergic decrease that take place offline early following learning^15^, possibly reflecting the role of neuronal excitation in the facilitation of synaptic plasticity processes^16–18^.

Here, we took advantage of the increased spatial, temporal, and spectral resolution of ultra-high field (7T) magnetic resonance spectroscopy (MRS) to address this discrepancy by measuring the concentrations of Glu and GABA in M1 before, immediately after, and 30 minutes following either brief motor memory reactivation, or repetitive motor skill practice. We demonstrate that brief motor memory reactivation induced significant offline improvements in skill performance measured on the following day, which were correlated to increased inhibitory tone in M1 immediately following the reactivation. Increased inhibition was expressed as both increased GABA levels as well as decreased Glu/GABA ratio (i.e., excitation-inhibition ratio/E-I ratio) in M1 immediately following the reactivation. Perhaps surprisingly, no inhibitory changes were linked with performance improvements following repetitive practice, suggesting that brief reactivation and repetitive practice differ fundamentally in terms of their early underlying offline neural processes. Alternatively, repetitive practice might engage additional competing mechanisms which override initial fast-acting inhibitory alterations. This latter interpretation could also explain the fact that both increased^14,19^ and decreased^20^ inhibitory function have been linked with memory improvements following reconsolidation. Equally surprising is the absence of a link between excitatory changes (either glutamatergic or E-I ratio) and the extent of offline learning in both the brief and repetitive practice groups, indicating that consolidation and reconsolidation also differ in the early mechanisms by which they lead to improvements in performance. Finally, our results emphasize the importance of an immediate (minutes) inhibitory response during a short time window, which is specific to brief reactivation-induced behavioral changes over longer timescales (days). Since the inhibitory changes were demonstrated during the initial offline period following the reactivation, when the reactivated memory is thought to be once again vulnerable to disruption or interference^6,7,11^, these fast inhibitory changes may reflect an important re-stabilizing mechanism, which is specific to brief reactivation of a memory trace. This possibility is further supported by previous findings demonstrating an important role of M1 in stabilizing new skill learning during the early stages of motor consolidation^21^.

## Results

### Motor Memory Reactivation Promotes Learning Gains

To address the question of the neurochemical basis of reactivation-induced motor learning, we used a task in which motor memory modifications have been previously demonstrated following its reactivation^10,11^. We implemented a 3-day repeated measures experimental design which follows the reactivation-reconsolidation framework^11,22^, which stems from investigations at the synaptic level^9^. Fifty-two young adults (age 27.4±3.8 years old, 24 females) underwent initial motor learning using a motor sequence learning task, in which they were asked to perform a five-digit tapping sequence over twelve practice blocks of 30 s each as fast and accurately as they could (see *Materials and Methods*)^2^. On the following day, participants performed either (i) a repetitive practice of twelve 30 s blocks, identical to the first day (i.e., Full-Practice group, n=17); (ii) A single 30 s block of reactivation of the motor skill memory (i.e., Reactivation group, n=18); Or (iii) no practice at all (i.e., Control group, n=17). The time duration of the reactivation followed works implementing similar procedures^2^, as well as previous findings suggesting that reactivation of less than 60 seconds is required to render a motor memory unstable again^6^. Finally, all participants were tested for their learning progress on the third consecutive day (see **Figure 1** for schematic illustration of the experimental design).

**Figure 1.**
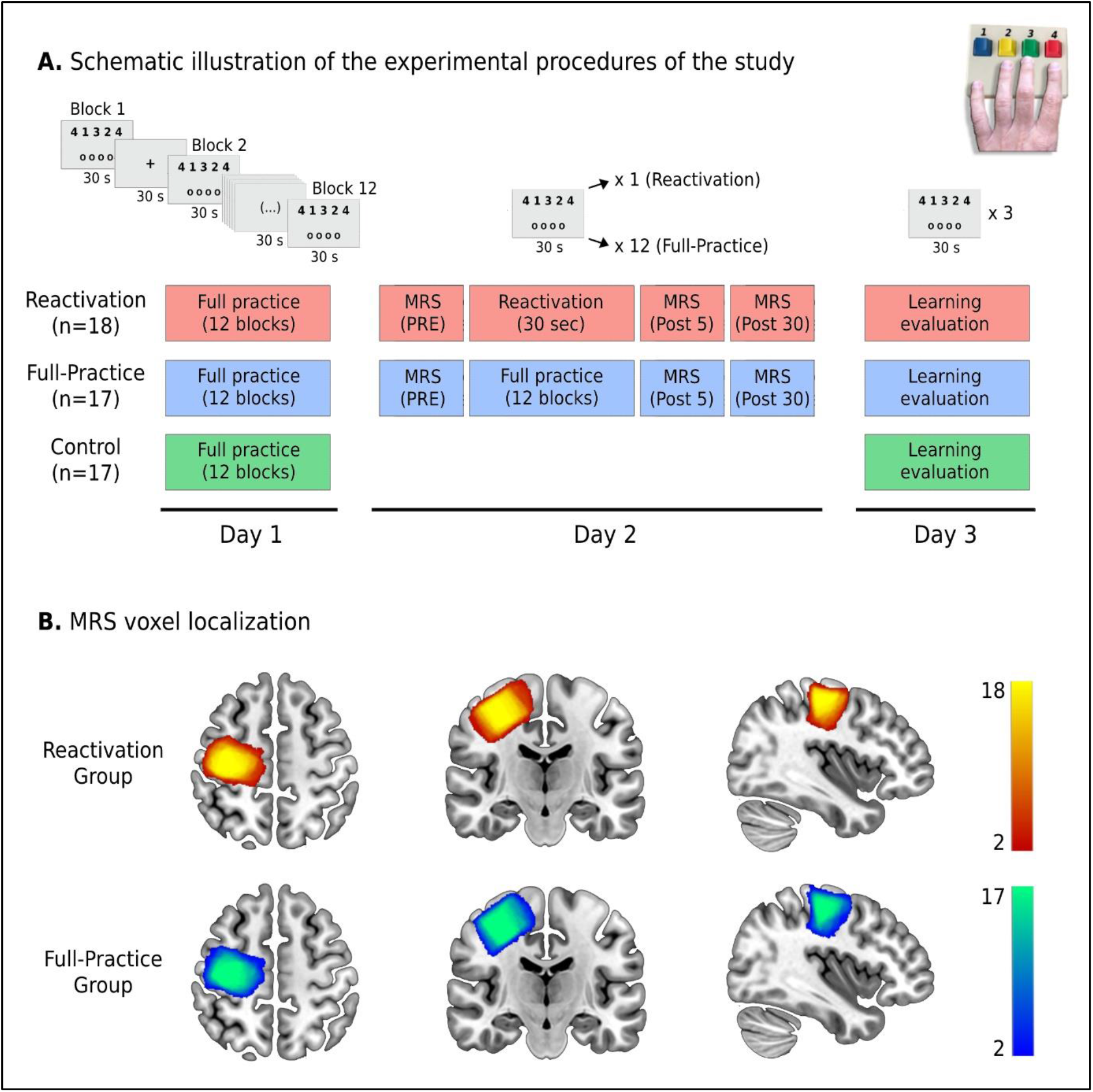
Experimental design and MRS voxel localization. The Reactivation and Full-Practice groups underwent three experimental sessions on three consecutive days which included initial motor skill learning on the first day, additional practice/motor memory reactivation on the second day and learning evaluation on the third day. The Control group underwent only initial learning and learning evaluation on the first and third days, respectively. The procedures on the first and third days were identical for all three groups. The sessions on the first two days were performed inside the MRI scanner, while learning evaluation was performed on a lab computer on the third day. MRS measurements were taken prior, immediately after, and 30 minutes following the practice/reactivation on the second day (A). MRS voxel localization: the MNI-transformed voxels within and across the Reactivation and Full-Practice groups are presented (B). Note the consistency in the placement of the MRS voxel on the right motor cortex within and across the groups.

We conducted a repeated measures linear mixed model analysis to test whether the motor learning paradigm elicited significant behavioral changes in skill performance within each of the three groups, comparing the performance on the last practice block of day 1 with the performance on the first block (i.e., the testing block) of the session on day 3. Skill performance was defined as the number of correct sequences performed in a given block, which is a common behavioral measure that combines both the speed and accuracy of the performed skill^2^. While we found a significant main effect of time on skill performance across the entire sample (F_1,49_ = 11.51, *p* = .001), participants also exhibited a significant time × group interaction (F_2,49_ = 8.44, *p* < .001). Within-group post-hoc analysis (corrected for multiple comparisons) revealed that participants in the Reactivation group demonstrated significant learning gains from day 1 to day 3 (β = 2.06, SE = 0.83, t_49_ = 2.46, *p* = .017), while the Control group demonstrated a non-significant decrease in performance (β = −1.00, SE = 0.85, t_49_ = −1.17, *p* = .249). The Full-Practice group demonstrated significant improvements in skill performance across the days as expected (β = 3.94, SE = 0.85, t_49_ = 4.59, *p* < .0001) (**Figure 2A**).

**Figure 2.**
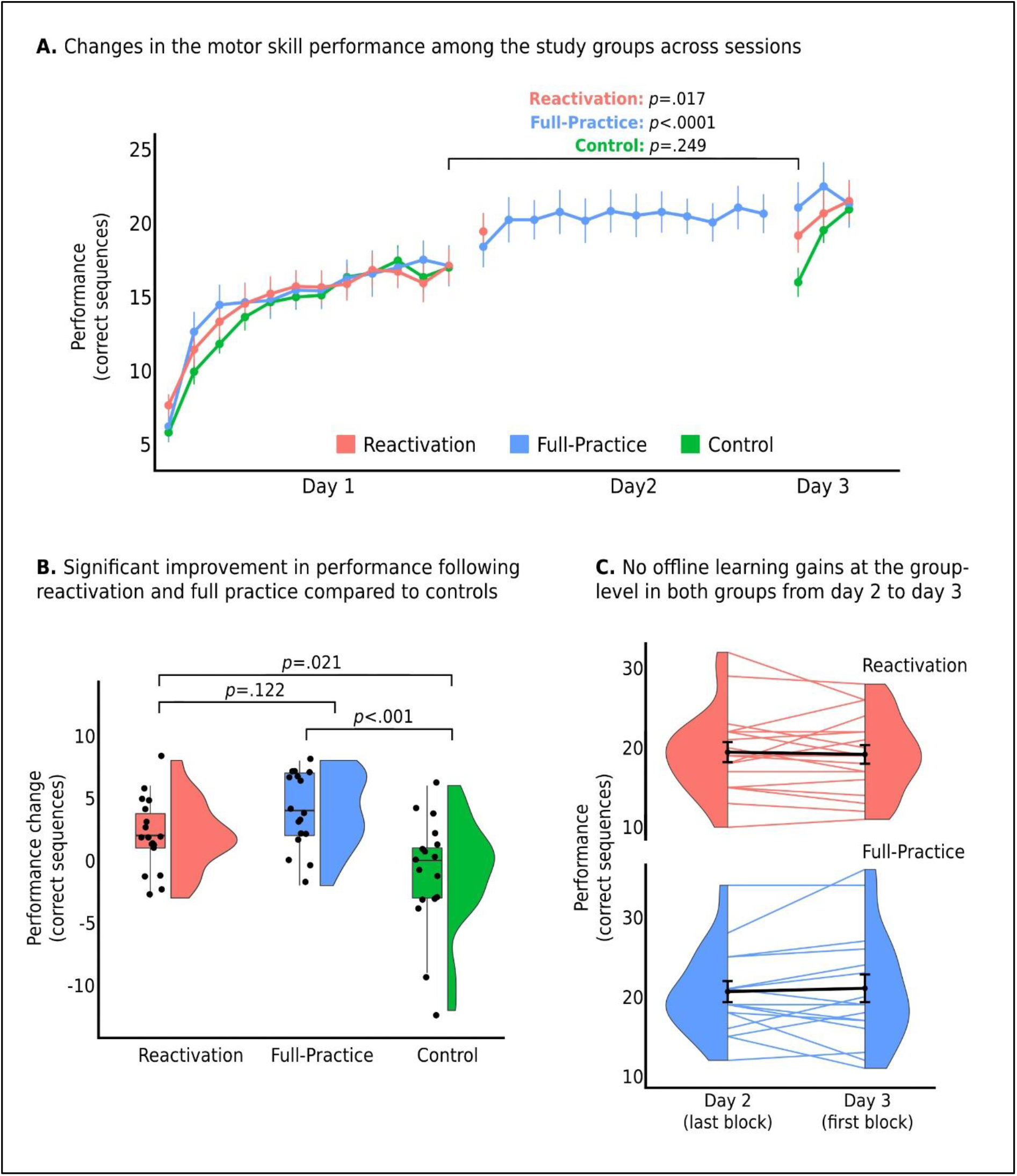
Differences in motor learning-induced behavioral changes within (A) and between (B) the study groups across sessions, demonstrating the efficacy of brief motor reactivation in promoting motor learning. In addition, although no group-level offline learning gains were found following the second practice session in both the Reactivation and Full-Practice groups, considerable variability was observed across participants on a between-subject basis (C, black lines and error bars represent each group’s mean±SEM).

In order to directly examine whether the groups exhibited different levels of learning gains, we used One-Way ANOVA to compare the extent of the learning gains observed across the days between the groups (i.e., day 3 performance minus day 1 performance). While all groups performed equally on the last block on the first day (*p* = .99 for all pairwise comparisons), both the Full-Practice group (β = 4.94, SE = 1.21, t_49_ = 4.07, *p* < .001) and the Reactivation group (β = 3.09, SE = 1.20, t_49_ = 2.55, *p* = .021) had significantly greater learning improvements compared to the control group across the three days (**Figure 2B**). In addition, although the average learning gains of the Reactivation group was lower than the observed in the Full-Practice groups, this difference was not statistically significant (β = −1.89, SE = 1.20, t_49_ = −1.58, *p* = .122). Furthermore, it has been previously proposed that the efficacy of motor memory reactivation in promoting further learning gains may depend on the quality of the execution of the reactivation^2^ (as quantified by the Continuity Score – see *Materials and methods*). However, we did not find evidence for a relationship between the Continuity Score and improvements in skill performance from day 1 to 3 among the participants in the Reactivation group (*r* = 0.16, *p* = .529).

Following the finding that both motor memory reactivation and repetitive practice promoted significant and indistinguishable motor learning gains, we aimed to examine whether the improvements in performance from day 1 to day 3 that were observed in both of the experimental groups were a result of learning gains that occurred online during the practice session on day 2, or following the session, i.e., as offline learning gains taking place between day 2 and day 3. To examine a possible offline learning effect, we conducted a repeated-measures linear mixed analysis comparing the performance on the last block on day 2 (i.e., the single block of the Reactivation group and the 12^th^ block of the Full-Practice group) and the testing block (i.e., the first block) on day 3. Surprisingly, we found no evidence for offline improvements between day 2 and 3 at the group-level, since no significant main effect for time between practice sessions (F_1,35_ = 0.02, *p* = .895) nor time × group interaction (F_1,35_ = 0.47, *p* = .498) were observed (**Figure 2C**).

The lack of group-level offline learning gains following a session of re-practicing a motor skill is in contrast to the significant overnight offline gains usually taking place following initial consolidation of explicit motor learning^11^, and which were also evident in the current study. The participants in both the Full-Practice and Reactivation groups demonstrated significant overnight offline learning gains following the first day of learning (F_1,33_ = 35.31, *p* < .001) with no significant differences between the groups (F_1,33_ = 2.90, *p* = .098) (**Figure 2A**). As with the overall learning gains across the three days, the Continuity Score was also not associated with the extent of offline learning gains following the motor memory reactivation among the participants in the Reactivation group (*r* = −0.16, *p* = .524). These findings in turn suggest that the behavioral improvements observed at the group-level in the two experimental groups from day 1 to day 3 may have resulted primarily from the offline learning gains during initial consolidation following the first day of practice, but not from offline gains between days 2 and 3. Additional group-level improvements between days were also driven by performance during the practice itself on the second day: When we examined the changes in performance across the 12 blocks of practice in the second day in the Full-Practice group (i.e., the difference in performance between the first and 12^th^ blocks) we found significant increases in skill performance (β = 1.32, SE = 0.41, *p* = .005) (**Figure 2A**), which in turn reflects the contribution of these within-session online learning gains to the overall improvements across the three days. Furthermore, when we included all twelve blocks in the repeated-measures model, only the difference between the 1^st^ and 2^nd^ blocks, and the 10^th^ and 11^th^ blocks were statistically significant (uncorrected for comparisons between all pairs of consecutive blocks). This in turn, may complement the effect seen after the single block in the Reactivation group and further support the importance of the first motor memory reactivation to the overall learning gains in a practice session, regardless of the total volume of that session.

### Increased Inhibition in M1 Following Reactivation Supports Overnight Offline Learning Gains

Despite no significant offline changes in performance at the group-level from day 2 to day 3 in any of the groups, there was considerable between-subject variability in the extent of performance changes that were evident overnight between those sessions (**Figure 2C**). We examined whether this between-subject variability could be explained by early neurochemical modifications of Glu and GABA in M1 following the reactivation/practice. Importantly, we have focused our examination to the M1 as this brain region has been widely implicated in the acquisition and consolidation of motor memories^12,14,23,24^. Furthermore, M1 has previously been linked to motor reconsolidation and the modification of motor memories following reactivation^10^.

We first ensured that the two groups were not different in their resting levels of either Glu (*p* = .191) or GABA (*p* = .971) before the practice on the second day. Then, we investigated whether there were significant changes in Glu or GABA immediately and 30 minutes following the second day practice compared to pre-practice levels (i.e., day 2 baseline levels) at the group-level. We found significant increases in Glu concentrations following practice (F_2,34.98_ = 4.97, *p* = .013), while GABA showed no significant change following the session (F_2,34.98_ = 1.00, *p* = .379). Post-hoc analysis revealed that Glu levels were significantly higher after 30 minutes (β = 0.27, SE = 0.09, t_36_.7 = 3.05, *p* = .013) but not immediately following practice (β = 0.10, SE = 0.06, t_37.1_ = 1.57, *p* = .125) compared to the resting levels at the beginning of the second day (**Figure 3A**). No time x group interactions were observed for either of the metabolites (*p* > 0.6), suggesting that the dynamics of either Glu or GABA following practice were not statistically different between the two distinct types of motor skill practice at the group-level. This in turn, may support the fact that the groups were also not significantly different in their extent of offline learning gains following the practice on day 2. However, as we highlighted above, there was a considerable variability in the extent of the offline learning gains between-subjects in both groups. Therefore, we next examined whether the extent of offline learning gains could be explained by the change in Glu or GABA concentrations, either immediately following the practice or after 30 minutes. We found that greater increases in GABA immediately following the motor memory reactivation were strongly associated with greater offline learning gains in the Reactivation group (*r* = 0.65, *p* = .004) (**Figure 3B**), with a moderate relationship observed with higher GABA increases after 30 minutes, although this relationship was not statistically significant (*r* = 0.40, *p* = .113) (**Figure 3C**). In contrast, changes in Glu were not related to the extent of offline learning gains in the Reactivation group, either immediately following the reactivation (*r* = −0.11, *p* = .653) or after 30 minutes (*r* = −0.15, *p* = .571) (**Figure 3B&C, respectively**). We found no relationships among the participants in the Full-Practice group between the extent of offline learning gains and post-practice modifications of either GABA (immediately: *r* = 0.23, *p* = .379; 30 minutes: *r* = 0.13, *p* = .608) or Glu (immediately: *r* = 0.21, *p* = .423; 30 minutes: *r* = 0.31, *p* = .229). The GABAergic-related inhibitory relationship with overnight behavioral improvements that was demonstrated in the Reactivation group was mirrored by the E-I ratio, with greater decreases in the E-I ratio (i.e., greater inhibition) being correlated to increased offline performance gains, both immediately following the motor memory reactivation (*r* = −0.65, *p* = .004) (**Figure 3B**) and after 30 minutes (*r* = −0.41, *p* = .103) (**Figure 3C**).

**Figure 3.**
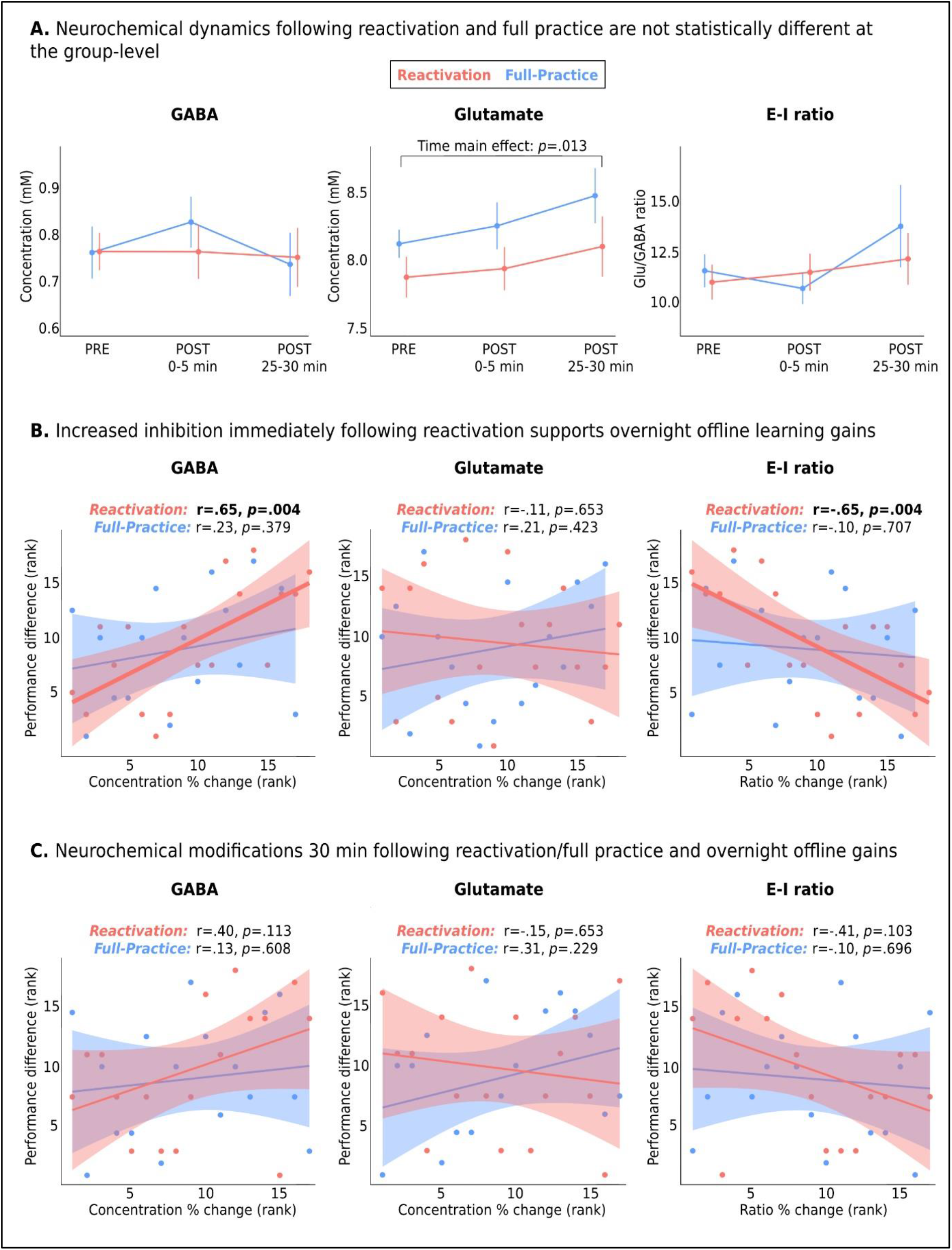
Neurochemical responses following the task on day 2. Glu levels significantly increased 30 minutes following the task with no significant differences between the groups, while no changes in GABA were observed at the group-level (A, mean+SEM are presented). Increased inhibitory tone immediately following the reactivation was correlated with overnight behavioral improvements (B). No significant relationship was found between neurochemical responses 30 minutes after the task and overnight changes in performance (C).

No significant correlations between the E-I ratio and offline performance gains were found in the Full-Practice group (immediately: *r* = −0.10, *p* = .707; 30 minutes: *r* = −0.25, *p* = .357). When we directly compared the strength of correlations between the two groups, we found the negative relationship between immediate E-I changes and offline learning gains to be significantly stronger in the Reactivation group compared to the Full-Practice group (*z* = 1.79, *p* = .037); A statistical trend was demonstrated for the relationship between offline gains and immediate changes in GABA (*z* = 1.46, *p* = .073).

We did not find a relationship between the resting (baseline) levels of Glu, GABA, or E-I ratio before the practice on the second day and performance in the first block of practice across all participants in both groups (*r* < 0.15, *p* > .21) or the Continuity Score in the Reactivation group (*r* < 0.15, *p* > .29). Statistical trends between greater online learning improvements during practice and lower pre-practice GABA in M1 (*r* = −0.38, *p* = .067), as well as higher E-I ratio (*r* = 0.36, *p* = .080), were observed among the participants in the Full-Practice group, in accordance with previous findings linking lower resting GABA levels in M1 with motor learning^24^.

## Discussion

The recently-reported phenomenon of reactivation-induced motor skill learning^2,25^ could potentially dramatically shorten the time required for motor skill learning or reacquisition. The aim of the current study was to reveal its underlying neurochemical mechanisms. We used a brief motor memory reactivation lasting 30 seconds that has previously been shown to render a consolidated skill memory labile again^6^, to examine how early post-reactivation modifications in GABAergic and glutamatergic concentrations in M1 may support overnight offline learning gains following the reactivation.

### Motor Memory Reactivation-Induced Skill Learning May Be More Robust Than Previously Thought

Our findings demonstrate that brief reactivation of a motor memory has the potential to promote continuous motor skill learning. These results are in accordance with previous findings highlighting the potential of brief procedural memory reactivation in promoting further offline learning gains^2,3,6^. Furthermore, this potential may be even more robust than previously thought: We found reactivation to promote significant learning gains not only within the reactivation group, but also compared to controls, regardless of the quality of the reactivations, which is in contrast to previous reports^2^. However, it should be noted that in the current study we examined the between-subject correlation between the reactivation quality (i.e., the Continuity Score) and behavioral improvements, while Herszage et al. (2021)^2^ based their analysis on a dichotomous measure by median-splitting the Reactivation group into higher and lower Continuity Score groups.

In the current study we implemented a 3-day experimental set-up based on the reconsolidation framework^22^. Therefore, the current study only included a single session of reactivation, primarily since our main goal was to examine the neurochemical changes taking place following a reactivation, and not the boundaries of behavioral improvements following brief reactivations. In contrast, Herszage et al. (2021)^2^ implemented more practice sessions in their study. More practice sessions may be needed to reveal to the true behavioural potential (and therefore also the limitations) of long-term practice that is based only on motor skill reactivations, compared to the more traditional prolonged repetitive practice. It is possible that the potential of reactivation to significantly improve a motor skill over time is limited to the first sessions of practice, when skill execution is yet to be perfected or automated, and that the superiority of repetitive training becomes more and more apparent as practice advanced through time.

### Inhibitory Mechanisms in M1 Support the Reconsolidation of Reactivated Motor Memory

Our findings show that early increases in GABA and commensurate decreases in E-I in M1 immediately after Reactivation – but not Full-Practice – are strongly correlated to overnight improvements in task performance (Figure 3B). What could be the biological rationale for increasing M1 motor cortical inhibitory tone following motor memory reactivation, and how does it promote longer, overnight changes in performance?

Previous research has suggested that the balance between excitation and inhibition plays a key role in determining the degree of cortical plasticity, and that increased excitation or disinhibition enhance neuronal responses to external stimuli and facilitate synaptic plasticity processes^16,17^. Furthermore, increased excitability, and not inhibition, has been demonstrated during memory consolidation following the encoding of new procedural skills in both humans and animals^14,26^, while facilitation of GABAergic transmission has been shown to impair initial memory formation^20^. In addition to the initial consolidation phase, a recent study also found a similar relationship following the reactivation of an existing visual perceptual memory^4^: In their work, Bang et al. (2018) used ^1^H-MRS to show that reactivating a perceptual memory was followed by an immediate increase in the E-I ratio at the group-level, although no significant changes were demonstrated for either Glu or GABA when examined separately. Their design, however, differed from our own considerably, making it difficult to draw parallels between the works: Their reactivation paradigm consisted of several practice blocks, as opposed to a single brief block of reactivation as implemented in our and previous works examining motor reactivation^2^; And they did not examine any reconsolidation-related behavioral changes following the reactivation, neither on the same day nor overnight, and consequently did not investigate the relationship between their reported neurochemical changes and further learning.

Mixed findings regarding the link between E-I and memory reconsolidation also arise from pharmacological studies. For example, low doses of clonazepam, a GABA_A_ receptor agonist, have shown to support overnight improvement of declarative memory when administered immediately following reactivation^19^. In contrast, other studies have suggested that administration of GABA_A_ agonists following reactivation may be detrimental for the reconsolidation of fear memories^20^. Interestingly, in the context of motor learning, a recent study in rodents has shown that while initial consolidation was associated with increased E-I ratio and suppressed GABAA inhibition in M1, these measures returned to pre-training baseline levels following the second day of practice as GABAergic transmission increased^14^.

We postulate that, while increased excitation or disinhibition seems to play a vital role during initial consolidation, increased inhibition plays an equally-important role during reconsolidation: That of re-stabilizing an existing memory trace upon its reactivation. When reactivating an existing memory trace, as in the current study, and by that rendering it unstable again and prone to adverse modifications, an immediate mechanism that re-stabilizes the memory trace may be necessary before plasticity-inducing processes take over to promote any additional learning. Newly acquired motor traces are prone to disruption during the first few hours following skill learning – for example, by disrupting M1 function immediately (but not several hours) after learning^21^ – which highlights the importance of the initial phase of motor memory consolidation in this process^11,27^. Previously consolidated motor memories can be equally destabilized^6,7^, and the initial offline period immediately following the reactivation may therefore be critical for the re-stabilization of the reactivated memory trace. Our results support this view, as we found immediate post-reactivation motor cortical inhibition to associate with overnight offline learning gains, that may be induced only after an unstable trace has been once again stabilized. The possibility that immediate motor cortical inhibition following reactivation may be necessary for the re-stabilization of the motor memory may be further supported by evidence from previous ^1^H-MRS studies in humans regarding the role of GABA and the E-I ratio in promoting plasticity versus stability^28^: Tamaki et al. (2020) found that changes in the E-I ratio during sleep following visual learning were associated with either facilitation or stabilization of the pre-sleep learning. Specifically, they showed that increased E-I ratio (i.e., greater excitation) in the visual cortex during the non-rapid eye movements (NREM) sleep stage was associated with greater post-sleep behavioral gains, but decreased E-I ratio (i.e., greater inhibition) during the REM sleep stage was associated with the stabilization of the pre-sleep learning. In addition, Shibata et al. (2017)^26^ have shown that over-learning a visual skill is associated with decreased E-I ratio and greater inhibitory tone in the visual cortex that strongly stabilizes the newly learned skill and protect it from interference. Furthermore, both Stagg et al. (2011)^24^ and Kim et al. (2014)^29^ demonstrated that smaller reduction in contralateral M1 GABA levels at rest (i.e., less disinhibition) induced by transcranial direct current stimulation (tDCS) was predictive of poorer motor learning gains. While those later findings are not directly related to reactivation or reconsolidation of existing motor memories, they further support the mechanistic link between the functionality of the GABAergic system and the potential to induce learning-related behavioral changes^24^. Lastly, our results are also in line with previous works demonstrating the importance of M1 processing during the early stages of initial motor memory consolidation, when the memory trace is still in an unstable state^15,21^.

### Inhibitory Changes Necessary for Reconsolidation Occur on Fast Timescales

In contrast to newly acquired memories, the re-stabilization of an existing memory trace may occur on faster timescales, as its representation in the brain has already been formed during initial consolidation. Indeed, it has been suggested that there is a limited time-window during which the memory is unstable and vulnerable following its reactivation^4,9^, and that this time period may be as short as tens of seconds in some instances^7^. Therefore, the initial offline period of reconsolidation immediately following reactivation is critical for the re-stabilization of the memory trace, and presumably also for offline learning gains to further develop across sleep^11,27^. These views are consistent with our findings, which show that increases to the inhibitory tone, presumably required for re-stabilizing reactivated memories happen on rapid timescales of minutes. Specifically, we have shown that an early inhibitory response immediately following the reactivation is associated with behavioral changes on longer timescales (Fig. 3B), and that this association weakens considerably just 30 minutes after reactivation (Fig. 3C).

Additional evidence for the timescales involved are obtained from the Full-Practice group, where increased inhibition and overnight improvements were not correlated between-subjects (Fig. 3C). This is consistent with the view that the re-stabilization of the motor memory has already been achieved following its initial retrieval in the first practice block of the full practice, and that the re-stabilization has already been completed during the practice session. Since all practice blocks in the Full-Practice condition presumably reactivated the same neural representation, this may not have interfered with the re-stabilization and may have even promoted a more efficient and effective process. This hypothesis is supported by the observation that the Full-Practice group demonstrated significant online learning gains already during the second day of practice. Therefore, the differential dependency on post-practice GABAergic responses observed between the two practice conditions in the current study may suggest that increased cortical inhibition may constitute a specific mechanistic response to brief memory reactivation, that may be required in order to re-stabilize the unstable memory trace during the limited time-window of reconsolidation. Furthermore, our results suggest that the extent of the response of this inhibitory mechanism may underpin individual differences in the potential of motor memory reactivation to induce further skill learning. Finally, it is important to note that this inhibitory effect does not seem to be a homeostatic consequence of task-related motor activation, since significant neurochemical changes were not observed immediately following either the Reactivation or the Full-Practice at the group-level.

## Conclusions

The current study proposes a potential neural mechanism at the micro-scale in the human brain for role of brief memory reactivation in promoting motor learning. Increased inhibition in the motor cortex immediately following motor skill reactivation, but not following traditional repetitive practice, supports the reconsolidation of the motor memory and further offline gains in skill performance that occur overnight. Our results emphasize the importance of the initial offline period following reactivation, and the role of an early neurochemical response in promoting behavioral changes on longer timescales. Furthermore, our findings highlight the important role of M1 in modifying existing motor memories following their reactivation. The results of the current study may have important clinical and rehabilitative implications for patients with impaired motor function, given the recent advances in non-invasive brain stimulation methods which can externally manipulate excitation and inhibition in the human brain^29,30^.

## Materials and methods

### Participants

Fifty-two healthy right-handed young adults, ages 18 to 40 (mean age 27.4±3.8 years, 24 females) participated in the current study. All participants provided written informed consent, approved by the Wolfson Medical Center Helsinki Committee (Holon, Israel), and the Institutional Review Board (IRB) of the Weizmann Institute of Science, Israel. Exclusions criteria included age below 18 years or above 40 years, musicians or video gamers (past or present), any neuro-psychiatric history (including medications), and participants who did not meet the safety guidelines of the 7T scanning policy.

### Experimental procedure

We conducted a between-group repeated measures experiment based on the 3-day framework of memory reconsolidation studies^22^. Each participant was randomly allocated into one of the three study groups. All participants performed the same procedures on the first day consisting of twelve 30 s blocks of motor sequence learning paradigm (see the *Motor learning paradigm* section below). On the next day, participants in the Full-Practice performed an additional 12-block practice session of the skill (identical to the first day) while participants in the Reactivation group performed only a single 30 s practice block. Lastly, on the third day, all participants from all three groups performed a final session of three 30 s blocks, with the first block comprising the “testing block” that reflected the learning outcome across the days^2^. Therefore, the Control group (n = 17) performed only the initial learning and final sessions. The experimental sessions on the first two days were conducted inside the MRI scanner, while the session on day 3 was performed on a lab computer. During the first and second days, participants underwent single-voxel MRS prior to and following the motor sequence task. Anatomical images were also acquired at the beginning of each scanning session.

### Motor learning paradigm

Participants performed an explicit motor sequence learning task in the MR scanner in which they were asked to repetitively tap a five-digit sequence (4-1-3-2-4) with their non-dominant left hand, as fast and accurate as possible^2^. Keypresses were performed on an MR-compatible response box with 4 computer-like pressing keys (Cedrus Corporation, Lumina LS-LINE model). The response box was placed near the left thigh, was adjusted for each participant’s arm length, and fixed to this position to prevent its movement during the scan. Keypress 1 was performed with the little finger, keypress 2 with the ring finger, keypress 3 with the middle finger and keypress 4 with the index finger. During each task block the five-digit sequence (4-1-3-2-4) was projected on the screen. Each task block also included four white circles presented on the screen, and each finger press was followed by a corresponding circle being filled for the time duration of that specific press. This procedure was implemented to provide the participant with online visual feedback (whether pressing on the desired key) and the experimenter with online information regarding the execution of the correct finger sequence. The visual circles did not provide error feedback, only information on the pressing finger at a given time. Consecutive task blocks were separated by a 30 s fixation block in which participants were asked to fixate on a black cross presented at the middle of a bright screen. The first day included a short pre-scan familiarity session with the task on a lab computer, in which only the experimenter performed a short version of the task with a control sequence, to prevent any learning effect in the participant prior to the actual task in the scanner. Therefore, the “real” sequence was only reveled to the participants when starting the learning paradigm. Importantly, all participants demonstrated at list one correct sequence pressing during the first learning block. Visual stimuli presentation and data collection were conducted using the Psychophysics Toolbox Version 3 (http://psychtoolbox.org/) implemented in MATLAB.

### MRI scanning procedures

#### MR data acquisition

The scanning sessions were performed on a 7T Terra scanner (Siemens-Healthineers, Erlangen, Germany) using a commercial single-channel transmit/32-channel receive head coil (NOVA Medical Inc., Wilmington, MA, USA), capable of maximum B1+ amplitude of 25 μT. Soft pads were used to hold each participant’s head in place to minimize head movement during the scanning sessions. An initial localizer and gradient echo field map were acquired for automated B0 shimming (Scan parameters for B_0_ mapping: TR/TE1/TE2=406/3.06/4.08 ms, *α*=25°, 1.9×1.9×2.0 mm resolution, TA=1:04 min). A high-resolution structural T1-weighted MP2RAGE (Magnetization Prepared 2 Rapid Acquisition Gradient Echoes) image was acquired for voxel placement and subsequent tissue segmentation (TR/TE/TI1/TI2=4460/2.19/1000/3200 ms, *α*1 = 4°/*α*2 = 4°, 1 mm^3^ isotropic voxels, TA=6:56 min). For the MRS acquisitions, a 2×2×2 cm^3^ spectroscopic voxel was placed over the hand knob region of the right primary motor cortex, based on neuroanatomical guidelines^31,32^ (**Figure 1B**). The voxel was shimmed using the automated B0 shimming capabilities of the in-house Visual Display Interface (VDI) libraries (The Weizmann Institute of Science, Israel, www.vdisoftware.net) in MATLAB 2020b (The Mathworks, Natick MA). The MRS acquisition was performed with a SemiLASER (sLASER) sequence (TR/TE=7000/80 ms, NEX=36, TA=4:58 min) previously optimized, and also validated for other brain regions as well ^33^. Functional MRI data were acquired using a multiband gradient-echo echo-planar imaging sequence. Scanning parameters were implemented according to the Human Connectome Project (HCP) 7T protocol^34^ (TR/TE=1000/22.2 ms, field of view=208×208 mm^2^, matrix size=130×130, voxel-size=1.6 mm^3^, 85 slices, multi-band/GRAPPA acceleration factor=5/2, bandwidth=1924 Hz/Px, flip angle=45°).

#### MRS analysis

MRS pre-processing was carried out using the VDI libraries. Coils were combined via signal-to-noise ratio (SNR) weighting, with weights computed from the reference water and noise scans, using a singular value decomposition algorithm. Spectra were aligned and phase-corrected relative to each other using a previously published robust iterative algorithm^35^. Global zero-order phase-correction was carried out based on the 3.0 ppm creatine peak in the summed spectra. No apodization or zero filling were employed. SPM12 (Wellcome Center for Human Neuroimaging, UCL, UK, http://www.fil.ion.ucl.ac.uk/spm) was used to segment the T1-weighted anatomical images into grey matter (GM), white matter (WM), and cerebrospinal fluid (CSF) images. Tissue fractions within the spectroscopic voxel were computed using VDI for subsequent use in absolute quantification and as a quality assurance metric. Metabolite quantification was carried out using LCModel^36^ version 6.3c, with a basis set containing 17 metabolites: aspartate (Asp), ascorbic acid (Asc), glycerophosphocholine (GPC), phosphocholine (PCh), creatine (Cr), phosphocreatine (PCr), GABA, glucose (Glc), glutamine (Gln), glutamate (Glu), myo-inositol (mI), lactate (Lac), N-acetylaspartate (NAA), N-acetylaspartylglutamate (NAAG), scyllo-inositol (Scyllo), glutathione (GSH), and taurine (Tau). Basis functions were simulated by solving the quantum mechanical Liouville equation using VDI, taking into account the full 3D spin profile and the actual pulse waveforms. Absolute quantification was carried out by correcting the metabolite concentrations provided by LCModel for tissue fractions estimated from the segmented images^37^, assuming a water concentration of 43.3 M in GM, 35.88 M in white matter (WM) and 5.556 M in cerebrospinal fluid (CSF). Relaxation correction assumed the same value of T2 for GM and WM. We also assumed no metabolites in CSF tissue fractions^38^. The long TR eliminated saturation effects and, consequently, no T1 corrections were required. In addition to the concentration, the relative Cramer Rao Lower Bound (%CRLB) for each metabolite was also obtained.

### Statistical analysis

Statistical analyses and visualizations were performed and constructed with MATLAB, and R 4.1.2. Group-level repeated measures were analyzed with linear-mixed models using the lme4 package implemented in R^39^. Each mixed effect model in the current study was examined as random intercept and random slope model, enabling the expression of different baseline levels but also difference in the extent of change for each evaluated measure across the participants. To this end, participants ID was used as the random effect and time as the fixed effect: *dependent variable* ~ *time* + (*Time* | *ID*). Post-hoc pairwise comparisons in which statistically significant effects were observed were corrected for multiple tests using the False Discovery Rate (FDR)^40,41^. Overall learning gains for the three groups were quantified as the difference in performance between the last block on day 1 and the first block on day 3. Performance was quantified as the total number of correct sequences tapped in each block. Offline learning gains between day 2 and day 3 were quantified as the difference in performance between the last block on day 2 and the first block on day 3. Online learning gains during the practice on day 2 within the Full-Practice group were quantified as the difference in performance between the first and last practice blocks. The Continuity Score was quantified as the maximal number of correct sequences tapped continuously without errors within the reactivation block^2^. For the MRS data, the mean concentration and standard deviation of each metabolite, as well as the mean and standard deviation of the CRLB were calculated for each of the time points. Quality assurance of the MRS data followed previously reported metrics: visual inspection for gross artifacts, such as lipid contamination and spurious echoes, metabolites concentrations that were three standard deviations away from the mean of all time-points measurements were excluded from further analyses, as well as water linewidths exceeding 15 Hz FWHM, and SNR of ≤30 from the LCModel output^12,33,42^. MRS data quality summary and LCModel fitting representation are presented in the **Supplementary Table 1** and **Supplementary Figures 1 & 2**. Within-group correlations were evaluated with Spearman’s correlation coefficient (two-tailed tests). Between-group comparison of correlation coefficients was performed with Fisher’s r-to-z transformation using the *“cocor”* package implemented in R.

