## Supplementary material for "Increased cortical inhibition immediately following brief motor memory reactivation supports reconsolidation and overnight offline learning gains"

### MRS quality metrics

The MRS quality metrics are presented in **Table S1** and **Figures S1 & S2**. No MRS measurements were excluded due to low SNR (i.e., <30), contaminated spectra, water linewidths (>15 Hz FWHM) (**Table S1**), or concentrations that were three standard deviations lower than the mean of all measurements across groups or time-points (values ranged from -2.44 to 2.67 SD) (**Figure S1** illustrates the distribution of GABA measurements across all participants and time-points). One participant from the Reactivation group did not have a POST-30 min MRS measurement due to time limitation in the scanner. No differences were found between the two groups in the quality metrics using two-tailed independent samples t-tests ($p$>.05). **Figure S2** demonstrates a representative spectrum acquired from one participant.

**Table S1.** MRS quality metrics (Mean±SD)

| | Reactivation group (n=18) | | Full-Practice group (n=17) | |
|---|---|---|---|---|
| | SNR | Water FWHM | SNR | Water FWHM |
| PRE | 60.72±5.8 | 10.51±1.1 | 61.47±5.6 | 10.62±1.1 |
| POST 0-5 min | 59.61±6.7 | 10.78±1.1 | 60.24±5.2 | 10.88±1.0 |
| POST 25-30 min | 59.76±6.6 | 10.86±1.0 | 58.59±6.1 | 11.29±1.1 |

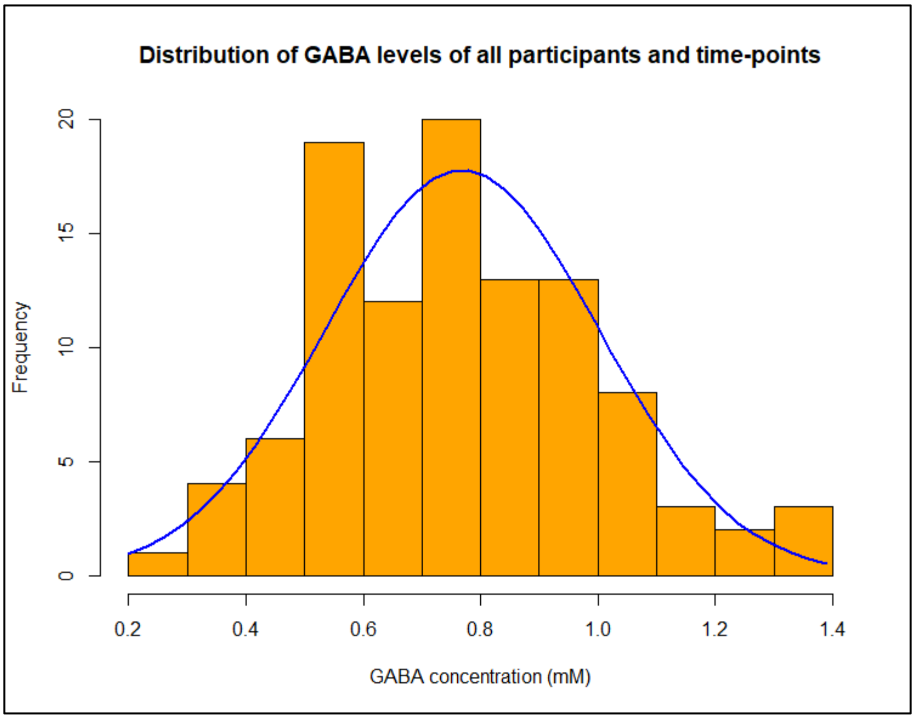


**Figure S1.** Histogram of GABA concentrations across all participants and time-points. A normal curve (in blue) is overlaid. Standardized values ranged from -2.44 to 2.67 SD.

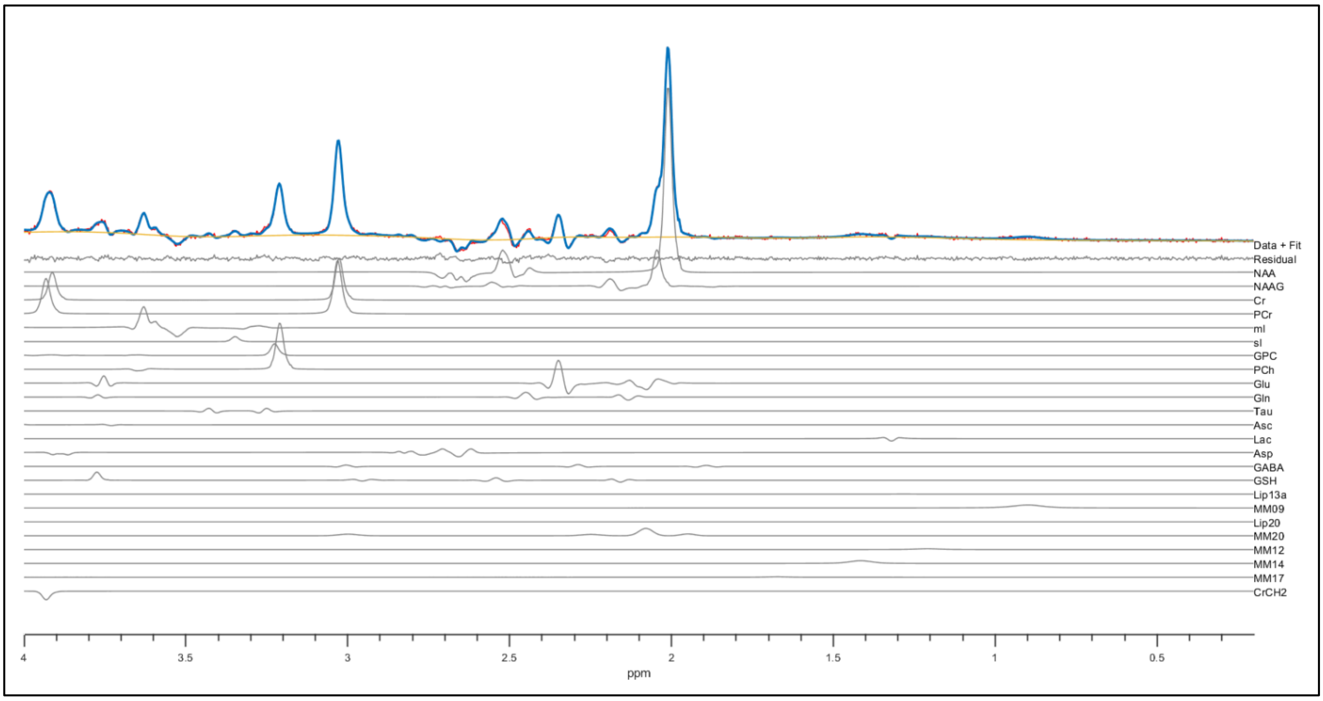


**Figure S2.** A representative spectrum acquisition from one participant including model fit. The data is presented with a red line, the fit with a blue line, and the baseline in yellow.
